# Deconvolution of membrane currents for low-crosstalk high-resolution membrane capacitance measurements

**DOI:** 10.1101/106641

**Authors:** Matej Hot’ka, Ivan Zahradník

## Abstract

Many cellular activities involving cell membrane manifest through changes in membrane capacitance. The accuracy and precision of cell membrane capacitance measurement is limited by the measuring system, which distorts membrane current responses used for analysis of impedance parameters. We developed a practical deconvolution procedure for reconstruction of the time course of membrane current and applied it to current responses recorded at step voltage changes by the whole-cell patch-clamp technique. In contrast to the recorded current responses, the reconstructed current responses could be exactly described by adequate impedance models of the recorded circuit. Deconvolution of membrane currents improved the performance of the square-wave method of high-resolution membrane capacitance recording by providing higher accuracy in a wider range of cell sizes and by eliminating cross-talk errors in parameter estimates. This allowed resolving instabilities in the recording conditions arising from parasitic capacitance and seal resistance variation. Complex tests on hardware models, on simulated data sets, and on living cells confirmed reliability of the deconvolution method for membrane capacitance recording. The aptitude of the method was demonstrated in isolated cardiac myocytes by recording spontaneous vesicular events, by discerning formation of a fusion pore, and by revealing artefacts due to unstable seal resistance.

## Introduction

Electrical phenomena originating at the cell surface can be used to characterize mechanisms of related cellular functions down to the single molecule level. The information content of the measurement is limited by the bandwidth and resolution of a recording method. Invention of the patch–clamp technique allowed high resolution measurements of electrical characteristics in most cells types. Nevertheless, some types of studies, including measurements of subtle changes in membrane capacitance associated with exo- and endocytotic membrane activity need special arrangements. The original high-resolution method of membrane capacitance measurement [1] has been subjected to several improvements [2,3]. These included the use additional equipment such as lock-in amplifiers [1,4,5], computer control hardware [6], or data acquisition boards with sophisticated software [7-9]. All these techniques provide excellent resolution but suffer from side effects of antialiasing filters [8-10] enhanced by instabilities under erratic recording conditions [2,3,11-13]. Sine-wave methods [1,10], based on single sine-wave voltage stimulation, are not well suited for cells with variable and nonlinear membrane resistance [2,11,12]. Dual sine-wave methods [5,7,12] solved these problems but at the cost of complexity and difficulty to optimize resolution for a given cell in experiment. Square-wave methods [8-10] showed a potential to outperform sine-wave methods due to their relatively simple implementation and low sensitivity to membrane resistance changes [2,8]. However, in measurements with a time constant below 200 μs, typical for cells of less than 40 μm in diameter, where the sine-wave methods excel, the square-wave methods start to suffer from errors instigated by an increased contribution of the parasitic capacitance, and by the low-pass antialiasing filter of patch-clamp amplifiers. Filtering suppresses the fast components of membrane current responses to step voltage changes, distorts their time course, and adds ringing at frequencies below the cut-off frequency [14]. This problem was originally minimized by blanking the first 70 μs of the current response, during which the filter effect is dominant [8-10]. As a result, however, a large part of the signal is lost, the amplitude of the remaining current is substantially decreased, and the number of data points used for the fitting procedure is reduced. Therefore, the square-wave methods had to be optimized for specific cells and experiments [8]; otherwise, the records of membrane capacitance are vulnerable to artefacts presenting as capacitance variances of unknown origin. These arise from instabilities in the recorded circuit (cross-talk), especially of the access resistance, but also of the membrane and/or seal resistance. Besides limiting precision and resolution, the artefacts downgrade reliability of the measurements.

The problem of low-pass filtering is common to membrane current measurements at step voltage pulses even when the voltage-clamp is fast. The high-frequency information in recorded membrane currents, active or passive, is suppressed by filtering. Therefore each improvement in fidelity of membrane current recordings would be beneficial in understanding of membrane processes.

Here we describe a method for reconstruction of the time course of passive membrane currents recorded by the whole-cell patch-clamp technique. In the core of the method is a deconvolution procedure that uses experimentally estimated transfer function of the recording system. The method was tested by hardware and software models as well as on isolated cardiac myocytes. We show that the deconvolution procedure cancels the distortions of the membrane current responses caused by low-pass filtering and improves the bandwidth of measurements. Application of the deconvolution method to high-resolution capacitance measurements allowed to eliminate cross-talk errors, and to treat difficult problems of parasitic capacitance interference and of the seal resistance instability.

## Materials and Methods

### Patch-clamp experiments

Experimental setup was built around inverted microscope Diaphot TMD (Nikon, Japan) placed on anti-vibration base Micro40 M6 (Halcyonics, Germany) and shielded by a Faraday cage. Cells were recorded in a 0.5 ml open air chamber with cover glass bottom (Warner Instruments, LLC, USA) and grounded with Ag/AgCl pellet electrode (Warner Instruments). Membrane current was recorded with an Axopatch 200B patch-clamp amplifier (Axon Instruments Inc., USA) using a glass micropipette fixed to a standard holder of the head-stage preamplifier CV-203BU (Axon Instruments Inc., USA) mounted on a 4-axis electrical micromanipulator MX 7600R (Siskiyou, USA). Patch pipettes were pulled from borosilicate glass capillaries (Sutter Instruments Co., USA) on Flaming/Brown Micropipette Puller Model P-97 (Sutter Instruments Co., USA). The pipette resistance was typically 2 - 4 MΩ. Pipette tips were coated with SYLGARD184 (Dow Corning, USA) to reduce stray pipette capacitance and its fluctuations. Formation of the seal between the patch pipette and the cell membrane was monitored by the pipette potential in cell-attached current-clamp mode [15]. A cell was accepted for measurements if the potential stabilized below -40 mV and withstood rupturing of the membrane patch. The final seal resistance was typically several tens of GΩs.

Ventricular myocytes of rat myocardia were isolated enzymatically [16] from young adult male or neonatal hearts of Han-Wistar rats (Dobra Voda, Slovak Republic). All anaesthetic and surgical procedures were carried out in accordance with the European directive 2010/63/EU, and were approved by the State Veterinary and Food Administration of the Slovak Republic (Ro-2821/09-221, Ro-354/16-221) and by the Ethical Committee of the Institute of Molecular Physiology and Genetics, Slovak Academy of Sciences.

Experiments on isolated cardiac myocytes were performed at room temperature of about 23°C. The standard bath solution contained (in mmol/l): 135 NaCl, 5.4 CsCl, 1 CaCl_2_, 5 MgCl_2_, 0.33 NaH_2_PO_4_, 10 HEPES, 0.01 TTX (pH 7.3; 300 mOsm). The composition of the pipette solution was (in mmol/l): 135 CsCH_3_SO_3_, 10 CsCl, 1 EGTA, 3 MgSO_4_, 3 Na_2_ATP, 0.05 cAMP, 10 HEPES (pH 7.1; 300 mOsm). Osmolarity of all solutions was verified using Osmomat 010 (Gonotec GmbH, Berlin, Germany).

### Impedance measurements and analysis

Whole cell impedance measurements were made using the patch-clamp amplifier in the whole-cell voltage clamp mode with a feedback resistor of 50 MΩ. The gain was set to 0.2 mV/pA to avoid amplifier saturation. The output current was low-pass filtered by the built-in 4-pole Bessel filter set to 10 kHz, digitized at 100 kHz by a 16-bit data acquisition system Digidata 1320A (Axon Instruments Inc.) and stored in proprietary Axon Binary Files (.abf) for off-line analysis.

Voltage stimuli were generated by Digidata 1320A 16-bit D/A converter. The amplitude of bipolar square-waves was as specified under individual figures. Stimulation period *T*_*S*_ was set in relation to the time constant *τ* of measurement as *T*_*S*_ = 12·*τ* (6·*τ* for each half-period [8]) and ranged from 20 to 0.6 ms (corresponding to stimulation frequency of 50 – 1,670 Hz). During impedance measurements, the holding potential was set to 0 mV. Experiments were controlled using pClamp 9 (Axon Instruments Inc.).

The recorded current responses to a square-wave voltage were processed by the deconvolution procedure or as specified. The parameters of current responses were estimated by fitting Eq. 1 to both half-periods of current responses *I_M_(t)* and averaged.

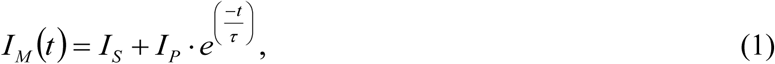

where *I*_*P*_ is the peak current, *I*_*S*_ is the steady current, and *τ* is the time constant. The parameters of the recorded cell circuit, approximated by membrane resistance *R*_*M*_, membrane capacitance *C*_*M*_, and access resistance *R*_*A*_ (Fig 1) were calculated from the estimated parameters of current responses (*I*_*P*_, *I*_*S*_, *τ*) using Eqs. 2A-C:

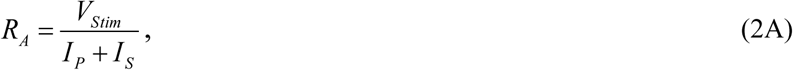

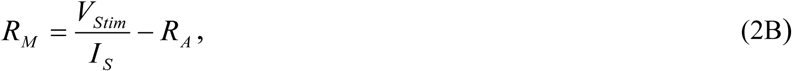

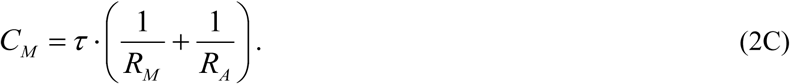

The analysis was performed off-line using routines written in MATLAB (v.2014b, Mathworks Inc., Natick, MA, USA) and implemented in the software MAT-MECAS (http://mat-mecas.sourceforge.net).

Standard deviation *σC*_*M*_ of the analysed time series of capacitance measurements, performed at the optimal stimulation frequency *f*_*S*_, was determined for the bandwidth *B* of 50 Hz. The number of averaged periods *m* was found using the relationship *m* = *f_S_ /B*. The theoretical limit of *C*_*M*_ resolution was calculated using Eq. 3 considering white thermal noise [8]:

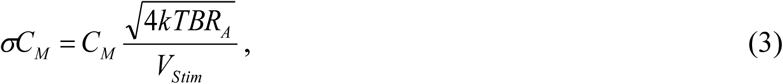

where *k* is the Boltzmann constant, *T* is the absolute temperature, and the remaining parameters were defined previously.

### Simulations and evaluation of errors

All simulations were performed in MATLAB (v.2014b, Mathworks).

To test the performance of the deconvolution procedure, we simulated 500 records of membrane current responses using Eqs. 4A-D and a seeding set of random triplets of circuit parameter values generated uniformly within ranges *C*_*M*_ (pF) ∈ [5, 200], *R*_*M*_ (GΩ) ∈ [0.02, 2], *R*_*A*_ (MΩ) ∈ [2, 20]:

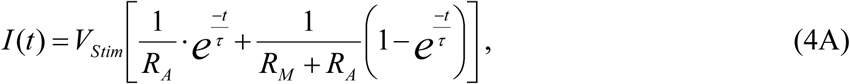

where

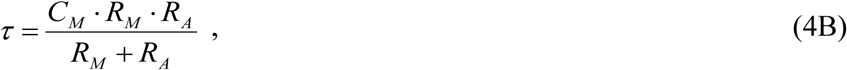

and the theoretical values or current parameters are:

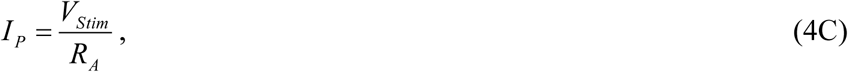

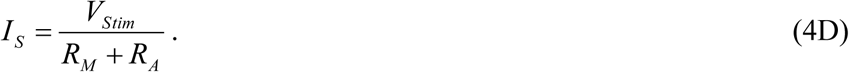

To simulate real recording conditions, the calculated membrane current responses were digitally filtered with the MATLAB function “filter”. The digital filter was an implementation of the analog 10 kHz 4-pole Bessel filter, using the method of invariance of impulse response “impinvar”, while the analog filter was emulated by the MATLAB function “besself”.

To evaluate the absolute error of estimate of circuit parameters, the simulated, digitized and filtered current responses were analysed in the same way as real records (using Eqs. 1 and 2A-C), and the difference between the seeded and the estimated parameter values was calculated.

For evaluation of the crosstalk error due to R_A_ variation, we simulated two periods of current responses for each random triplet of parameter values, so that the R_A_ value of the second period was increased by 1 MΩ. The simulated current responses were analysed in the same way as real records using Eqs. 1 and 2A-C. Crosstalk errors were evaluated as the difference between the estimated parameters of the two sets of simulated current responses.

To assess the effect of *R*_*A*_ variation on the resolution of membrane capacitance measurements, the current responses were calculated for a cell circuit with *C*_*M*_ = 50 pF, *R*_*M*_ = 200 MΩ, and *R*_*A*_ values fluctuating randomly around the mean of 5 MΩ. The resolution was evaluated as standard deviations *σC*_*M*_ estimated at the corresponding standard deviations *σR*_*A*_ (22.6, 45.2, 90.4, 135.6, 180.8, 361.7, and 452.1 kΩ), at the bandwidth normalized to 50 Hz. The dependence of circuit parameter estimates on the value of seal resistance *R*_*Seal*_ was simulated using Eq. 5,

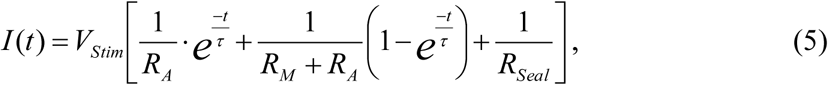

which differs from Eq. 4A by the last term that includes the *R*_*Seal*_ value. The current responses for a cell with *C*_*M*_ = 140 pF, *R*_*M*_ = 500 MΩ, and *R*_*A*_ = 4 MΩ were stimulated with *V*_*Stim*_ amplitude of ±10 mV and a period of 6*τ*. The seal resistance *R*_*Seal*_ was incremented by 100 MΩ period by period from 1 to 150 GΩ.

## Results

### The deconvolution procedure

The circuit in Fig 1A corresponds to the whole-cell patch-clamp recording configuration of a standard cell represented by the membrane capacitance, *C*_*M*_, and the membrane resistance, *R*_*M*_, connected to the recording amplifier via access resistance, *R*_*A*_. Resistance of the seal between the recording microelectrode and the cell membrane, *R*_*Seal*_, should be high, and capacitance of the recording microelectrode, *C*_*P*_, should be low to maximize the bandwidth of membrane current measurement.

**Figure 1.**
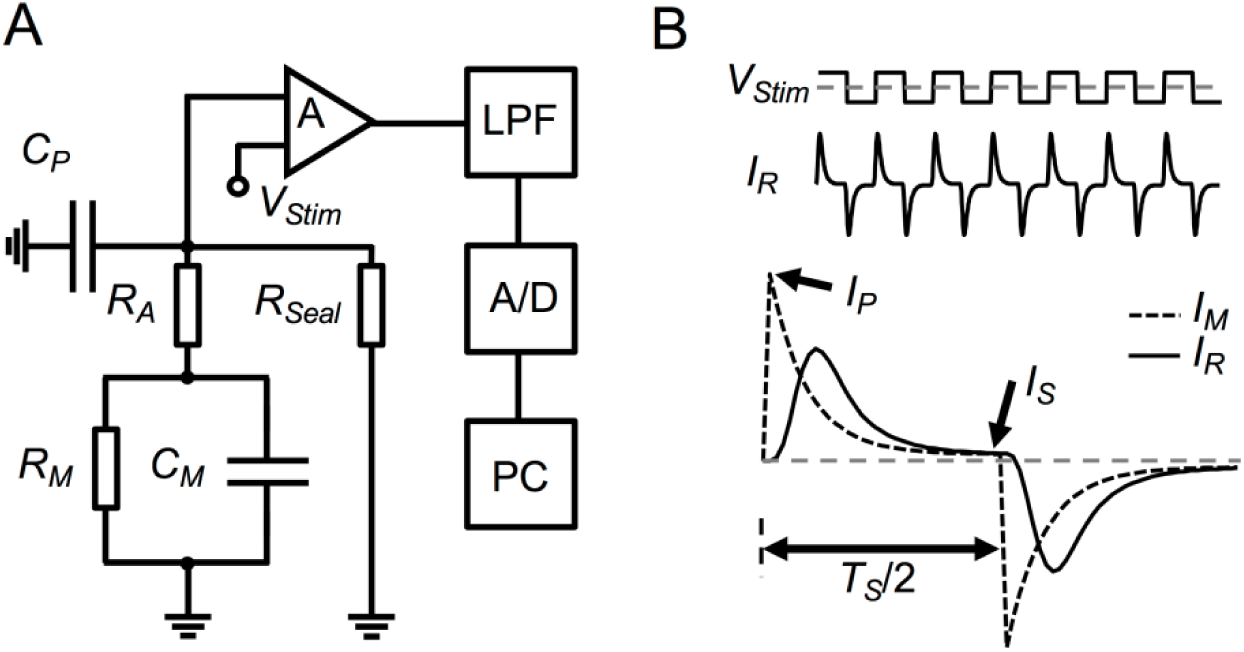
Patch-clamp recording and square-wave stimulation schemes. **(A)** Equivalent circuit of the whole-cell patch-clamp recording configuration. *C*_*M*_ - membrane capacitance. *R*_*M*_ - membrane resistance. *R*_*A*_ - access resistance. *R*_*Seal*_ - resistance of the seal between the recording microelectrode and the cell membrane. *C*_*P*_ - parasitic capacitance of the recording microelectrode. *V*_*Stim*_ - stimulation voltage applied to the measured circuit. A - patch-clamp amplifier. LPF - low-pass filter. A/D - digitizer. PC - computer. **(B)** Membrane current responses to square-wave voltage stimulation and a single-period current response. *V*_*Stim*_ - stimulation voltage. *I*_*R*_ - recorded current responses. *T*_*S*_ – period duration. *I*_*M*_ - the real membrane current response. *I*_*P*_ - peak membrane current. *I*_*S*_ - steady membrane current. Grey dash line - zero current level.

In theory, if the standard cell is voltage-clamped homogenously, a square-wave voltage stimulus, *V*_*Stim*_, induces a membrane current response, *I*_*M*_(*t*), of a peak amplitude, *I*_*P*_, that decreases mono-exponentially to a steady value, *I*_*S*_, approaching the DC limit at *t*_*∞*_ (Fig 1B). The membrane current response can be described by Eq. 1. In practice, however, the recording patch-clamp amplifier with a low-pass filter at the output distorts the current response and the recorded current does not comply with Eq. 1. Therefore, to minimize the errors in estimates of membrane current parameters, the recorded current should be processed appropriately before analysis.

The recorded current, *I*_*R*_(*t*), carries the measured current together with information about the signal path [17], due to convolution of the input current with the impulse response, *h*(*t*), of the signal path:

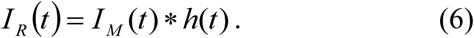

The original measured signal can be recovered by a deconvolution process, if the impulse response of the recording system is known. For this purpose we recorded the current impulse response of the recording system (Fig 2A). The voltage impulse was represented by a 10-μs 150-mV voltage stimulus (1 sample generated by the digitizer set to 100 kHz). When delivered to the amplifier configured with a resistor between the input and the ground, the impulse generated a complex current response (R+C response), which contained the resistive component (R response) combined with the capacitive component (C response) generated by the parasitic capacitance and the inner circuitry of the amplifier. These two capacitive components are always present in the current recording path. Their contribution to the signal should be eliminated to obtain the correct impulse response, *h*(*t*). To this end, the voltage impulse was delivered to the amplifier with the same resistor connected to the input but disconnected from the ground. In this arrangement, the current response contained only the capacitive component (C response). The R+C and C responses, each of 20 ms duration, were recorded 4096 times and averaged to supress the noise. The average C response was subtracted from the average R+C response and the resulting average R-response was transformed by the fast Fourier transform algorithm (FFT) to the frequency domain and normalized to 0 - 1 range. This operation yielded the system frequency characteristic *H*_*S*_ in 0.05 – 50 kHz bandwidth (Fig 2B). A special attention was given to proper grounding of the set-up to eliminate high frequency interference signals, because their presence in the *H*_*S*_ function caused oscillations in the reconstructed currents.

In practice, *H*_*S*_ is applicable to reconstruction of current responses measured with the same sampling and filtering frequency. For best results in cell experiments, the charging current of parasitic capacitance of the recording microelectrode should be properly cancelled either by using the circuitry of the amplifier or by subtraction of pre-recorded capacitive current responses, both in the cell-attached configuration before breaking the patch membrane.

Reconstruction of membrane current responses distorted by the recording path was performed in three steps. First, the recorded current response was transformed to the frequency domain by FFT. Then, the deconvolution was performed in the frequency domain by dividing the recorded current response by *H*_*S*_. Finally, the result was transformed to the time domain by inverse FFT, which yielded the reconstructed current response (Fig 2C).

The reconstructed current responses to square voltage stimuli followed exponential time course and could be approximated by Eq. 1. Consequently, the estimated current parameters corresponded to parameters of the theoretical circuit current and could be used for calculation of the impedance parameters of the recorded circuit using Eqs. 2A–D.

**Figure 2.**
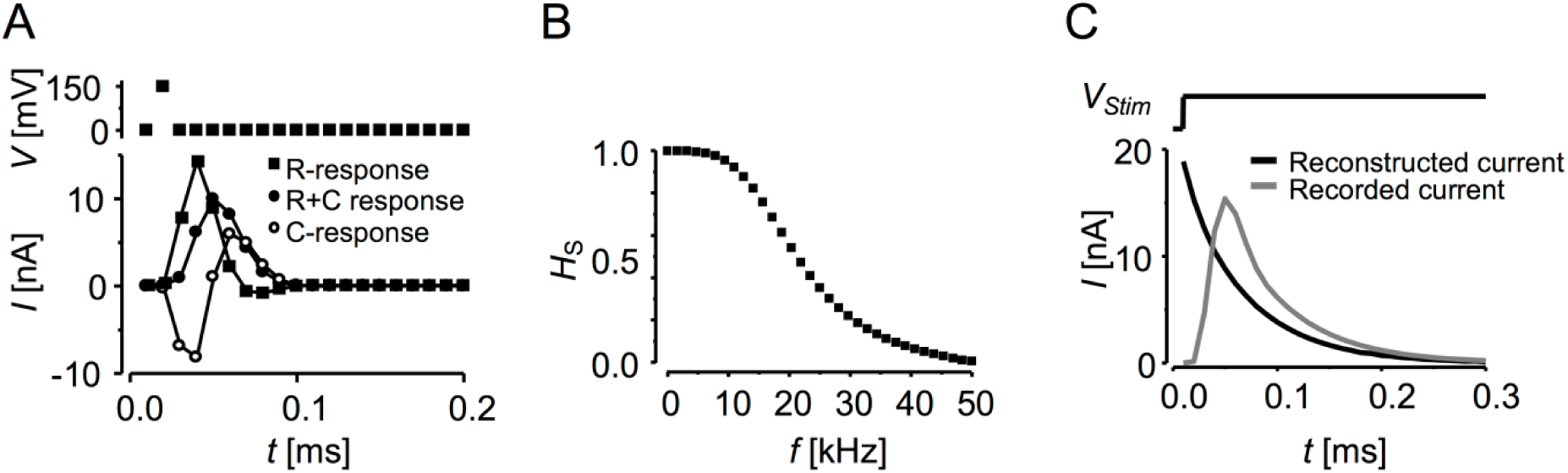
The deconvolution procedure. **(A)** Sampled voltage stimulus and current responses. *Upper panel*: the voltage impulse. *Lower panel*: the sampled current responses. (●) – the average current response recorded with the 4.7 MΩ resistor connected between the amplifier input and the ground (R+C response); (○) - the average current response recorded with the 4.7 MΩ resistor connected to the amplifier input and disconnected from the ground (C-response); the average current responses were obtained from 4096 records. (■) - the impulse current response (R-response) obtained by subtracting the C-response from the R+C response. **(B)** The characteristic of the signal path in the frequency domain, *H*_*S*_, obtained by FFT of the R-response. **(C)** An example of the recorded low-pass filtered current response (grey line) and of the current response reconstructed by deconvolution procedure (black line) obtained with the hardware model cell (*C*_*M*_ = 10 pF, *R*_*M*_ = 500 MΩ, *R*_*A*_ = 4.7 MΩ) to *V*_*Stim*_ = 80 mV. The parasitic capacitance was compensated by the amplifier compensation circuitry.

The whole processing and analysis procedure for deconvolution of membrane current records was written into an interactive software package MAT-MECAS in MATLAB v. 2014b (see: http://mat-mecas.sourceforge.net), which was used also in this study. The software reads records of current responses to square-wave voltages, saved in an appropriate format (the Axon Binary File abf; Axon Instruments Inc., or a text file), and creates records of the time course of *C*_*M*_, *R*_*M*_, *R*_*A*_, and other specified parameters in xlsx format. MAT-MECAS also provides special tools for secondary analyses, including identification of events and artefacts in *C*_*M*_ records. A record of 1000 current responses, each of 100 samples per period, was analysed in about 60 s at full temporal resolution (1 kHz), or in 3 s if the bandwidth was reduced to 50 Hz.

### Accuracy and resolution

The errors arising from the deconvolution procedure itself were assessed in the simulated set of filtered current responses generated with the set of random triplets of circuit parameters, as described in Materials and Methods. The simulated responses were processed by the deconvolution procedure and fitted with Eq. 1. Differences between the estimated parameters of current responses and the ones calculated for the given triplet of circuit parameters are plotted in Fig 3A. The deconvolution procedure provided reconstructed currents almost identical to the original ones, since the differences *∆τ*, *∆I*_*S*_, and *∆I*_*P*_ were negligible at any combination of the circuit parameter values.

**Figure 3.**
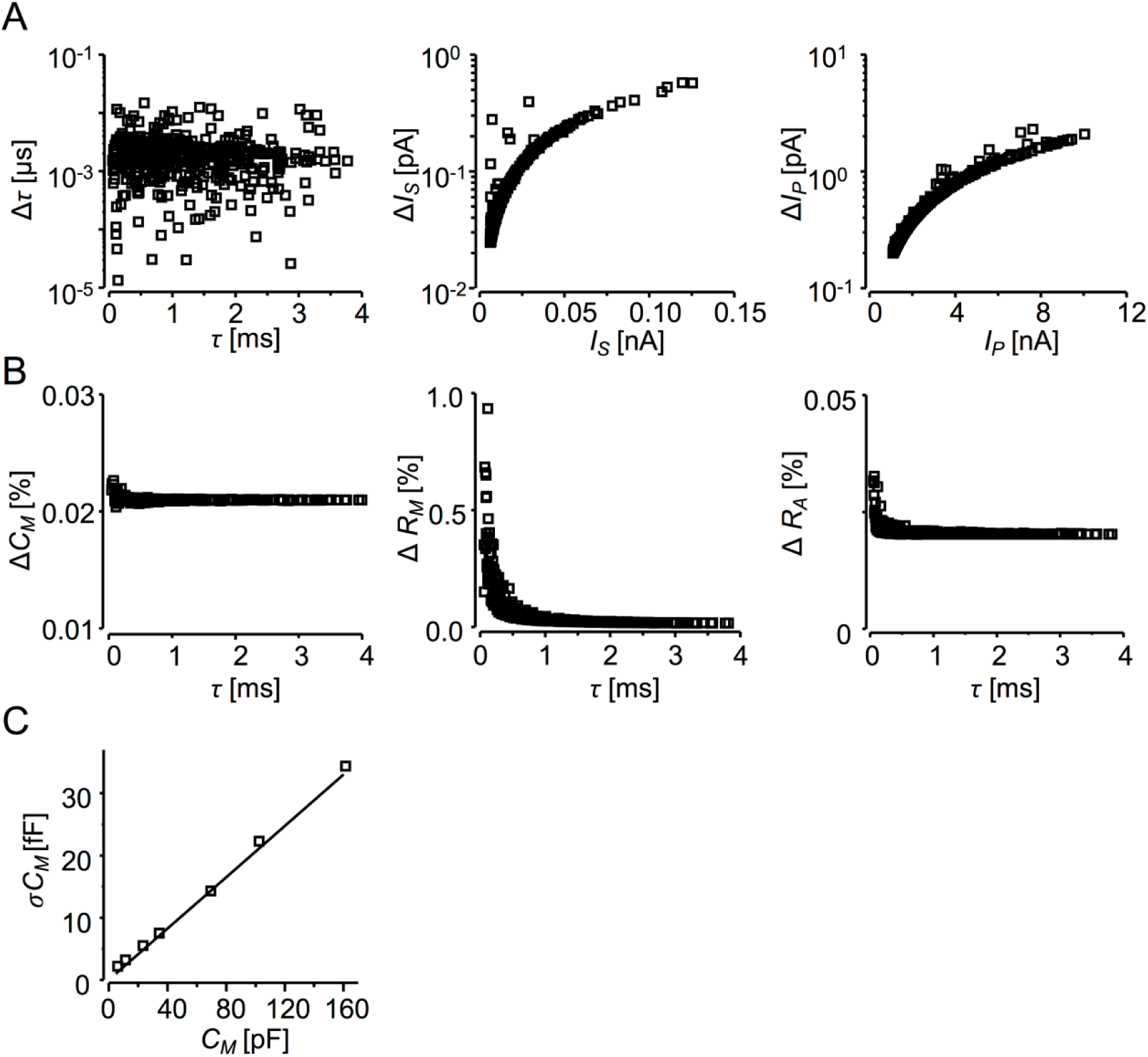
Accuracy and resolution tests of the deconvolution method. (**A**) Absolute errors of the estimated parameters of simulated, filtered, and reconstructed current responses (∆*τ*, ∆*I*_*S*_, ∆*I*_*P*_) plotted against their respective theoretical values. The errors were calculated as differences between the best fit parameter values of Eq. 1 to simulated, filtered, and reconstructed current responses and the theoretical parameter values calculated by Eq. 4B-D for the set of random triplets of cell parameters (see Materials and Methods). (**B**) Relative errors of the circuit parameters estimates (∆*C*_*M*_, ∆*R*_*M*_, ∆*R*_*A*_), determined from the simulated, filtered, and reconstructed current responses (Eq. 4), plotted against calculated *τ* values (Eq. 4B) for the respective random triplets of circuit parameters (the same dataset as in **A**). (**C**) Standard deviations of membrane capacitance σ*C*_*M*_ of the hardware model cells plotted against *C*_*M*_ values wired in the models (4.7, 10, 22, 33, 68, 101, or 160 pF). In all hardware models *R*_*A*_ was 4.7 MΩ and *R*_*M*_ was 500 MΩ. Square-wave ±10 mV, 12*τ* periods. The theoretical limit of *C*_*M*_ resolution (black line) was calculated by Eq. 3 considering thermal noise. Bandwidth 50 Hz.

Differences between the circuit parameter values used for simulation and those estimated by the deconvolution procedure are shown in Fig 3B. We plotted the differences as the percentage of the parameter value in the triplet against the corresponding *τ* values calculated for each triplet using Eq. 4B. The time constant of current response *τ* is a convenient reference parameter since it incorporates all circuit parameters. It also determines the reliability of the fitting procedure since, due to the constant sampling interval, the number of sampled data points is proportional to *τ*. The relative errors in *C*_*M*_ were under 0.02% and virtually independent of *τ* at any combination of circuit parameter values. The relative error in *R*_*M*_ estimates was less than 0.1% for *τ* > 500 μs and increased to about 1% for *τ* = 50 μs. The relative error in *R*_*A*_ estimates was very low, under 0.03% for *τ* > 100 μs at any combination of parameter values.

The resolution limit of the deconvolution procedure for membrane capacitance measurements was estimated on hardware models made of different capacitors combined with membrane and access resistors (Fig 3C). The deconvolution procedure performed near the physical limit of *C*_*M*_ resolution given by thermal noise (Eq. 3), even for very small cells (*C*_*M*_ < 20 pF).

### Cross-talk error

In measurements on real cells, all circuit parameters may vary in a rather wide range. Therefore, the parameter cross-talk is a crucial issue of any high-resolution membrane capacitance recording method. Of special consideration is the access resistance, which solely defines *I*_*P*_, contributes substantially to *τ*, and may vary by up to several MΩ on short as well as long time scales. At the same time, the cell membrane resistance *R*_*M*_ may change from a few GΩ to a few tens of MΩ due to experimental interventions or membrane impedance nonlinearities. On the other hand, *R*_*M*_ contributes substantially to the time constant only when comparable to *R*_*A*_, which is not a standard situation.

Propensity of the deconvolution method for cross-talk error was tested using the same set of parameter triplets as that used for testing the accuracy, to which siblings differing in *R*_*A*_ by 1MΩ were generated. Fig 4A shows results of analysis of the simulated current responses related to *τ*, through which the interdependence of parameters arises. Except for the very short time constants, the cross-talk of a 1 MΩ change in *R*_*A*_ to *C*_*M*_ and *R*_*M*_ was negligible. The differences in *C*_*M*_ estimates were typically much less than 1 fF, while the estimates of *R*_*A*_ changes were almost exactly 1 MΩ (0.9995 MΩ).

The role of cross-talk of *R*_*M*_ variation to other parameters was studied in additional simulations. These revealed that even very large variation of membrane resistance does not present a problem, since the method managed to keep errors of *C*_*M*_ estimation below 0.2 fF when *R*_*M*_ was changed by 100 MΩ in the 0.02 – 2 GΩ range (not shown).

Additional assessment of the crosstalk error was performed using the built-in compensation circuitry of the recording amplifier [8]. Recordings shown in Fig 4B were made with the input of the amplifier open and with the membrane capacitance compensation set to about 25 or 130 pF to simulate a small or a large cell, respectively. The series resistance compensation was set to about 5.5 MΩ in both cases. During recording, the series resistance compensation was changed to about ~6.5 MΩ and back to emulate fast and large *R*_*A*_ variation. Absence of visible correlated changes in the *C*_*M*_ and *R*_*A*_ traces confirms results of numerical simulations.

In real cell experiments, *R*_*A*_ can vary broadly at various time scales. In the presence of cross-talk, variation of *R*_*A*_ would transpire to *C*_*M*_ recordings, increase the variability of *C*_*M*_ estimates and decrease the resolution of *C*_*M*_ measurements. To assess the effect of cross-talk on *C*_*M*_ resolution, we simulated traces of current responses of a typical cell with *R*_*A*_ fluctuating randomly and compared the standard deviation of *R*_*A*_ with the standard deviation of *C*_*M*_ (Fig 4C). With the use of the deconvolution procedure, variance of *C*_*M*_ due to cross-talk from *R*_*A*_ was three orders below the variance that would result from thermal noise. If the current responses were corrected only by blanking the first 70 μs and fitting the remaining record of the current responses, the resulting crosstalk error increased fluctuations of membrane capacitance estimates above the level expected for thermal noise substantially for *R*_*A*_ changes exceeding 20 kΩ.

**Figure 4.**
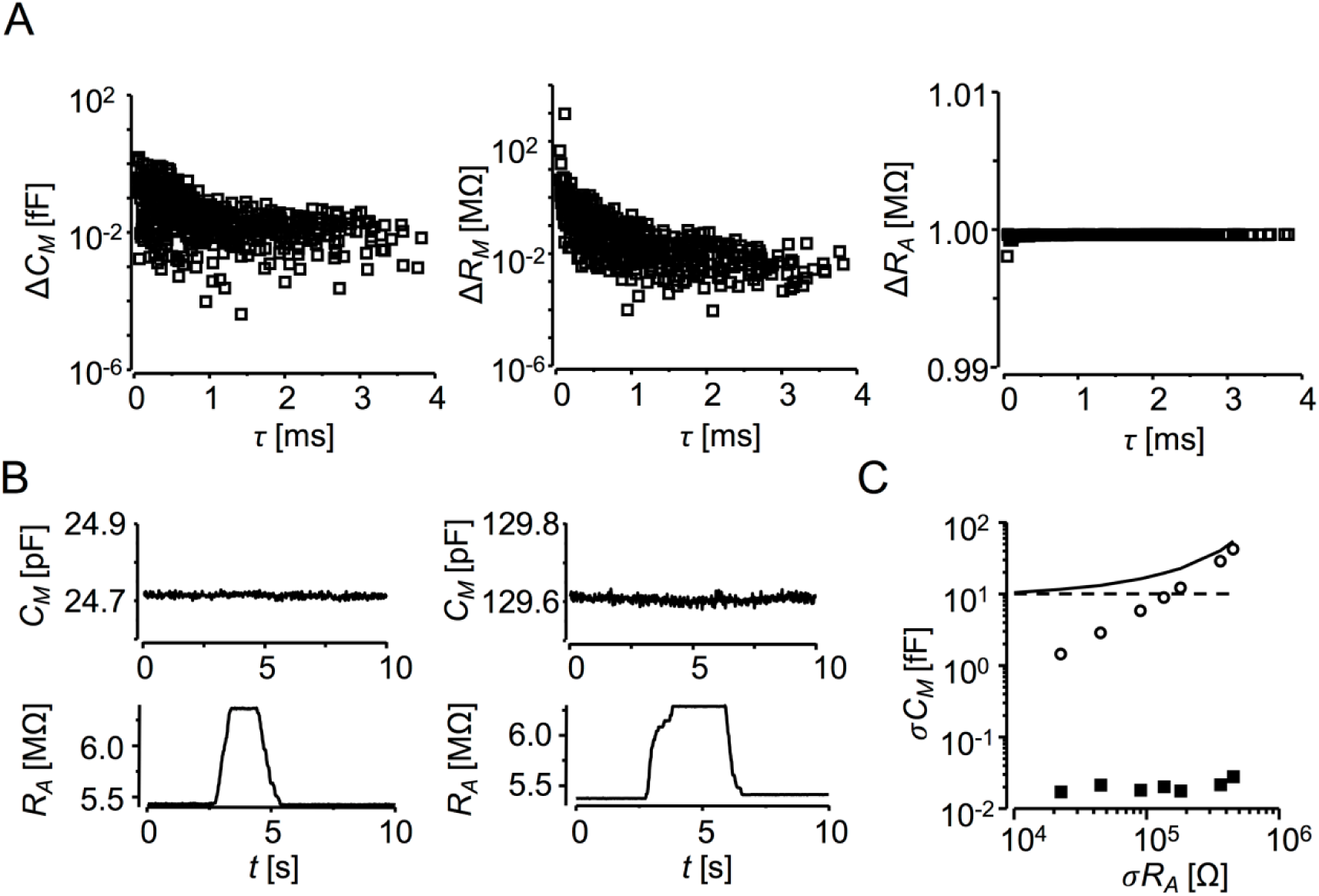
Tests of the deconvolution procedure for cross-talk errors. **(A)** Analysis of the simulated, filtered, and reconstructed current records. The differences in estimates of *C*_*M*_, *R*_*M*_, and *R*_*A*_ before and after increasing the *R*_*A*_ value by 1 MΩ is plotted against the range of pertinent *τ* values. **(B)** Analysis of current records obtained with the use of compensation settings of the recording amplifier. Square-wave ±40 mV, 10 ms periods, bandwidth 50 Hz. The *C*_*M*_ value was set to emulate either a small cell (left panels, *τ* = 136 μs) or a large cell (right panels, *τ* = 713μs). *Upper panels*: time series of *C*_*M*_ estimates. *Lower panels*: time series of *R*_*A*_ estimates. **(C)** Contribution of the *R*_*A*_ to *C*_*M*_ crosstalk error to the overall *C*_*M*_ resolution. A set of filtered current responses was simulated for a cell of *C*_*M*_ = 50 pF, *R*_*M*_ = 200 MΩ, and *R*_*A*_ varying randomly with the same mean of 5 MΩ but different values of standard deviation (500 stimulation periods for each σ*R*_*A*_). The standard deviation σ*C*_*M*_ in the *C*_*M*_ time series was estimated for 50 Hz bandwidth. Dashed line – the thermal noise level calculated by Eq. 3. (■) – the values of σ*C*_*M*_ in the time series of *C*_*M*_ estimates from current responses reconstructed by the deconvolution method. (○) – the values of σ*C*_*M*_ in time series of *C*_*M*_ estimates from current responses corrected by blanking the first 70 μs and extrapolating the best-fit function (Eq. 1) to the time of the peak current response. Black line – the sum of noise level and the data shown as circles. Notably, in the case of the deconvolution procedure, the fluctuations in *C*_*M*_ caused by fluctuations of *R*_*A*_ were by 3 orders of magnitude below the thermal noise level.

### Effect of parasitic capacitance

The parasitic capacitance, or the stray capacitance, of a recording microelectrode, *C*_*P*_, connects to ground in parallel to cell membrane. Since charging of *C*_*P*_ by the stimulus is not attenuated by the cell impedance, it is charged faster than the cell membrane. Nevertheless, the charging current *I*_*Cp*_ is reconstructed by the deconvolution procedure as a part of the recorded current (Fig 5A, upper panel). Presence of *I*_*Cp*_ caused dumped oscillation on the current response at the onset of a step voltage change, which arose due to Gibbs phenomenon [17], and may cause an error in estimation of the peak current amplitude.

**Figure 5.**
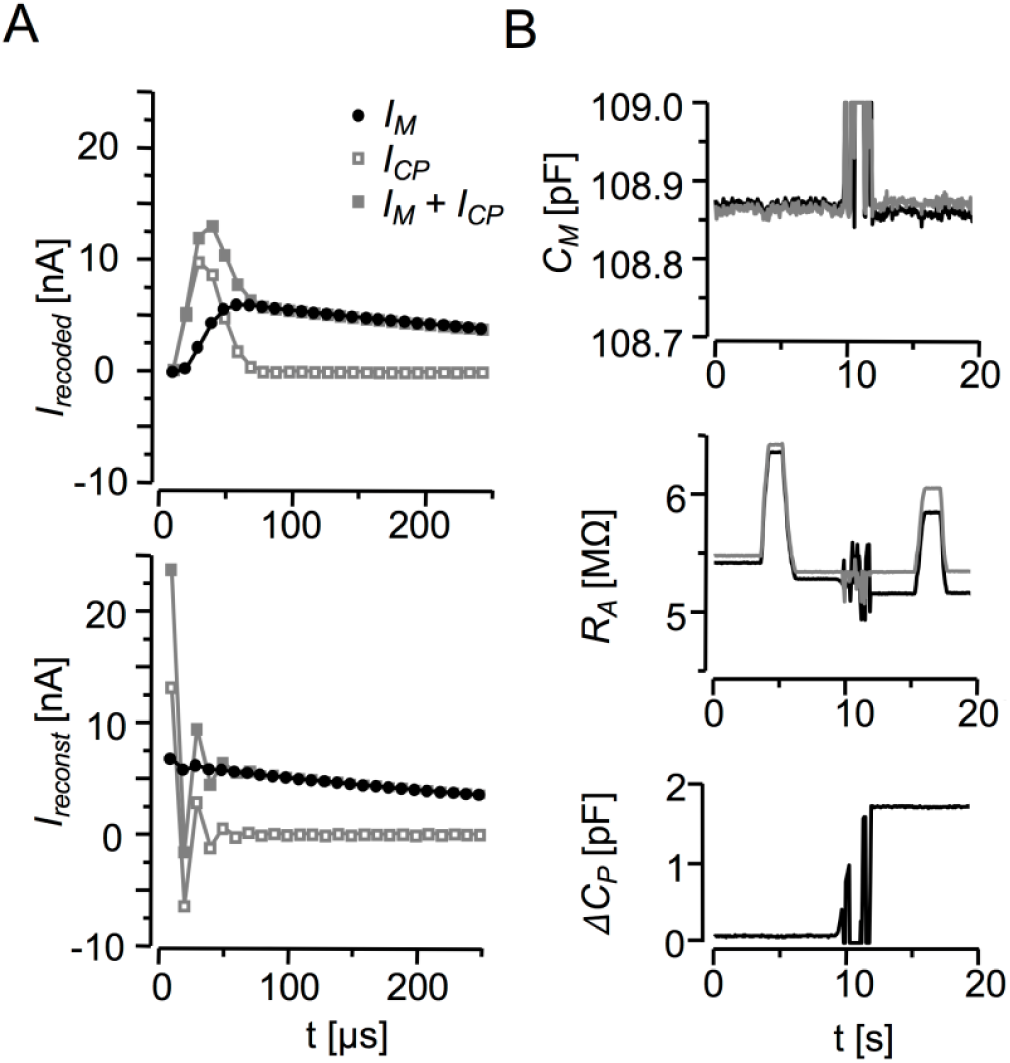
The effect of parasitic capacitance on estimates of impedance parameters. **(A)** Onset of current responses to step voltage changes of the measuring amplifier with shielded open input and with the compensation circuitry set to emulate cell circuit parameters. *Upper panel*: (●) – the current response (*I*_*M*_) recorded for *R*_*A*_ set to 5.5 MΩ, *C*_*M*_ set to 109 pF, and the input capacitance compensated. (**□**) – the current response record (*I*_*CP*_) of a parasitic capacitance emulated with decompensated fast capacitance setting of the amplifier (*R*_*A*_ and *C*_*M*_ set to zero). (■) – the current response (*I*_*M*_+*I*_*CP*_) recorded with combined settings of the amplifier as in *I*_*M*_ and *I*_*CP*_ records. *Lower panel*: the current responses corresponding to records in the upper panel reconstructed with the deconvolution procedure. (■) – reconstructed (*I*_*M*_+*I*_*CP*_) current response. (**□**) – reconstructed (*I*_*CP*_) current response. (●) – reconstructed (*I*_*M*_) current response with compensated parasitic capacitance; note the negligible oscillation. **(B)** Effects of changes of parasitic capacitance (see lower panel) on estimates of *C*_*M*_ and *R*_*A*_ of a cell emulated with the compensation circuitry of recording amplifier. Black traces – time series of parameter estimates of current responses reconstructed by the deconvolution procedure. Grey traces – time series of parameter estimates of current responses reconstructed with the deconvolution procedure applied to current responses with the first 70 μs blanked; note the absence of *C*_*P*_ crosstalk in *C*_*M*_ trace. The flickering artefacts are caused by handling of the fast capacitance compensation potentiometer. The change of parasitic capacitance Δ*C*_*P*_ was evaluated from the integral of the onset of recorded current response [9].

To estimate the error due to the presence of *I*_*Cp*_ in current records, we emulated changes of parasitic capacitance using compensation circuitry of the recording amplifier (Fig 5B). When the standard deconvolution procedure was applied to recorded current responses, the relatively large change in *C*_*P*_ setting by 1.8 pF (Fig 5B, lower panel) caused a *C*_*P*_ cross-talk both to *C*_*M*_ estimates (Fig 5B, upper panel, black trace) and to *R*_*A*_ estimates (Fig 5B, middle panel, black trace). Nevertheless, the *C*_*P*_ crosstalk was fully supressed when the deconvolution procedure was applied to the current responses with the first 70 μs blanked to omit the oscillations due to uncompensated parasitic capacitance (Fig 5B, upper and middle panel, grey traces). Comparison of black and grey traces in Fig 5B indicates that partially uncompensated stable *C*_*P*_ affects the estimates of cell impedance parameters minimally, but its temporal variation would cause a moderate crosstalk artefact in *R*_*A*_ and a minor one in *C*_*M*_. Moreover, the access resistance setting was also shortly changed (by about 0.9 and 0.7 MΩ, respectively) at each *C*_*P*_ level in this experiment (Fig 5B, middle panel). Again, as in previous tests, crosstalk of *R*_*A*_ to *C*_*M*_ records was below the resolution of the *C*_*M*_ measurement, indicating that oscillations of the current due to uncompensated parasitic capacitance have negligible effect on *R*_*A*_ crosstalk to *C*_*M*_.

To overcome the problem of *C*_*P*_ interference, the pipette capacitance should be recorded in the cell-attached configuration before going to whole-cell configuration, and subtracted from the current records. This procedure removes the oscillations from the deconvolved current responses. In practice, stabilization and minimization of the pipette stray capacitance by coating the patch-pipette close to its tip with a hydrophobic resin and compensation of the stray capacitance with built-in amplifier circuitry was sufficient for suppression of *C*_*P*_ effects in typical experiments (not shown). Nevertheless, if large changes of the bath level in the recording chamber had occurred, for instance during perfusion experiments, blanking of the first samples of current responses before the deconvolution analysis could be applied (Fig 5B, grey traces).

### Tests on real cells

The quality of the deconvolution procedure was tested on cardiac myocytes. In Fig 6A, a small isolated cardiomyocyte of neonatal rat heart displays spontaneous stepwise increases in membrane capacitance of about 10 to 50 fF in amplitude without any changes in membrane conductance and with a stable access resistance during the recording period. At 50 Hz bandwidth, the standard deviation of signal noise in the *C*_*M*_ trace was 2.6 fF.

The recording in Fig 6B (isolated cardiomyocyte of adult rat heart) shows capacitive changes under less stable recording conditions when the access resistance increased gradually by 550 kΩ and the membrane conductance contained single channel-like activity of up to 80 pS conductance. Nevertheless, the deconvolution method neatly resolved stepwise capacitive events of 40 to 80 fF in the *C*_*M*_ trace without any cross-talk from *R*_*M*_ and *R*_*A*_ variation. At 50 Hz bandwidth, the standard deviation of signal noise in the *C*_*M*_ trace was 11.5 fF.

**Figure 6.**
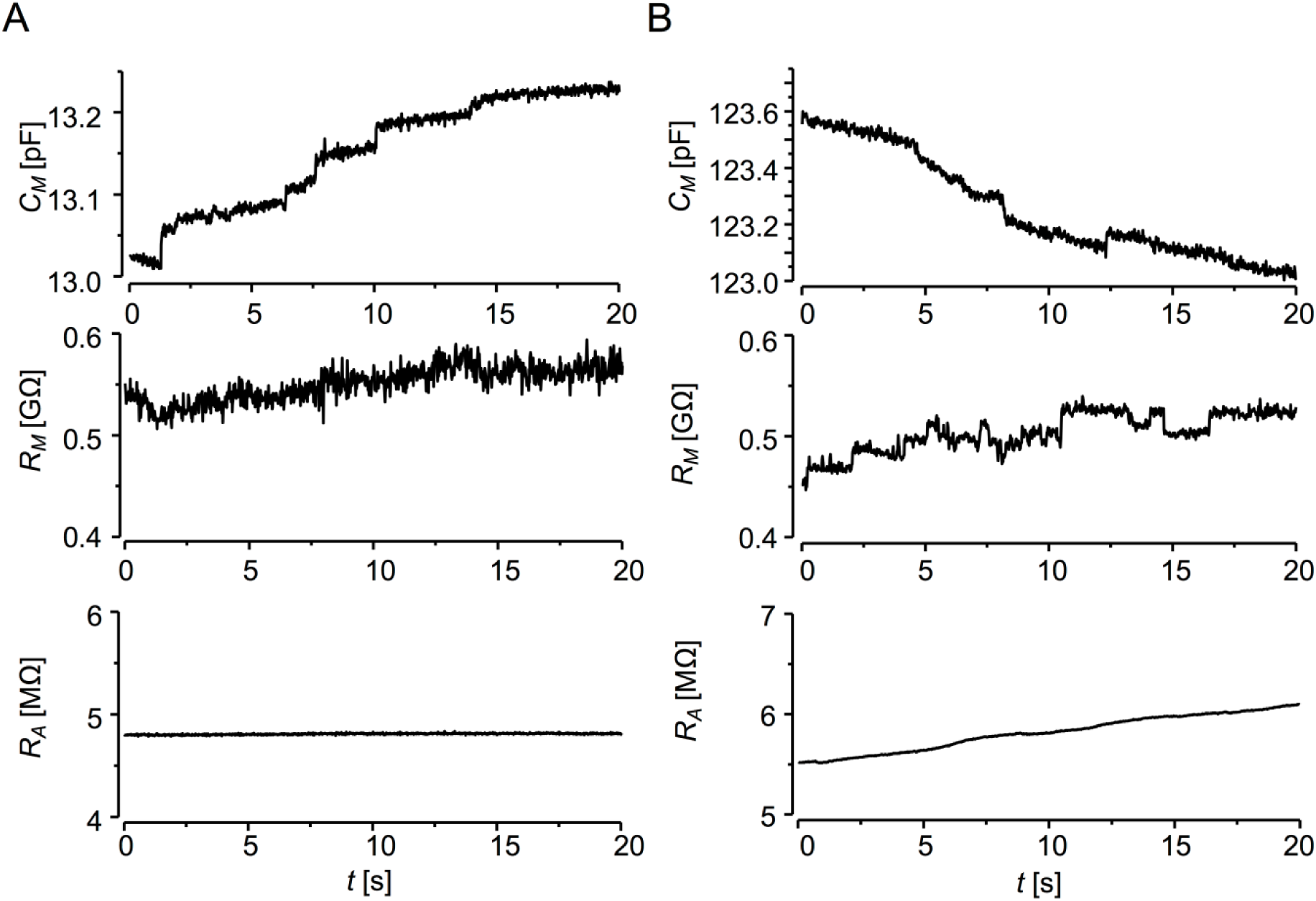
Tests of the deconvolution procedure on isolated cardiac myocytes. Note the absence of correlated changes in *C*_*M*_, *R*_*M*_ and *R*_*A*_ records indicating no visible cross-talk. (**A**) Recording on a neonatal rat heart cardiomyocyte. Square-wave ±40 mV, 1.25 ms period, bandwidth 50 Hz. (**B**) Recording on an adult rat heart cardiomyocyte. Square-wave ±40 mV, 10 ms period, bandwidth 50 Hz.

The square-wave voltage method can resolve the transient pore opening associated with fusion of an intracellular vesicle with the plasmalemma [8]. In Fig 7 we document the ability of the deconvolution method to resolve membrane capacitance events connected with prolonged opening of a fusion pore. The capacitance event started with a gradual, not stepwise, increase at about 1.7 s of the trace time. At the same time, *R*_*M*_ trace showed a small negatively correlated change, despite the fact that the deconvolution procedure is free of cross-talk errors. Together with the accompanying increased error of the fit of current responses (Fig 7, bottom panel) this indicated that the three-element impedance model (Eq. 1) was inappropriate for description of the current. We employed a five-element model for vesicle fusion [8] described by Eq. 7:

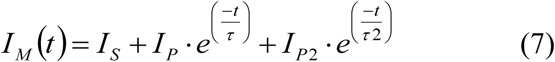

where *τ2* = *C*_*v*_/*G*_*p*_, *C*_*v*_ is the capacitance of the fusing vesicle equal to the amplitude of the capacitance increase, *G*_*p*_ is the variable conductance of the pore, and *I*_*P2*_ is the increase of the peak current amplitude due to fusion event (see [8] for details of the fitting procedure). The resulting fusion pore conductance increased from 0.1 to 0.5 nS within 250 ms and then jumped to an incalculable value, indicating that the pore opened wide and the fusion process had been completed.

**Figure 7.**
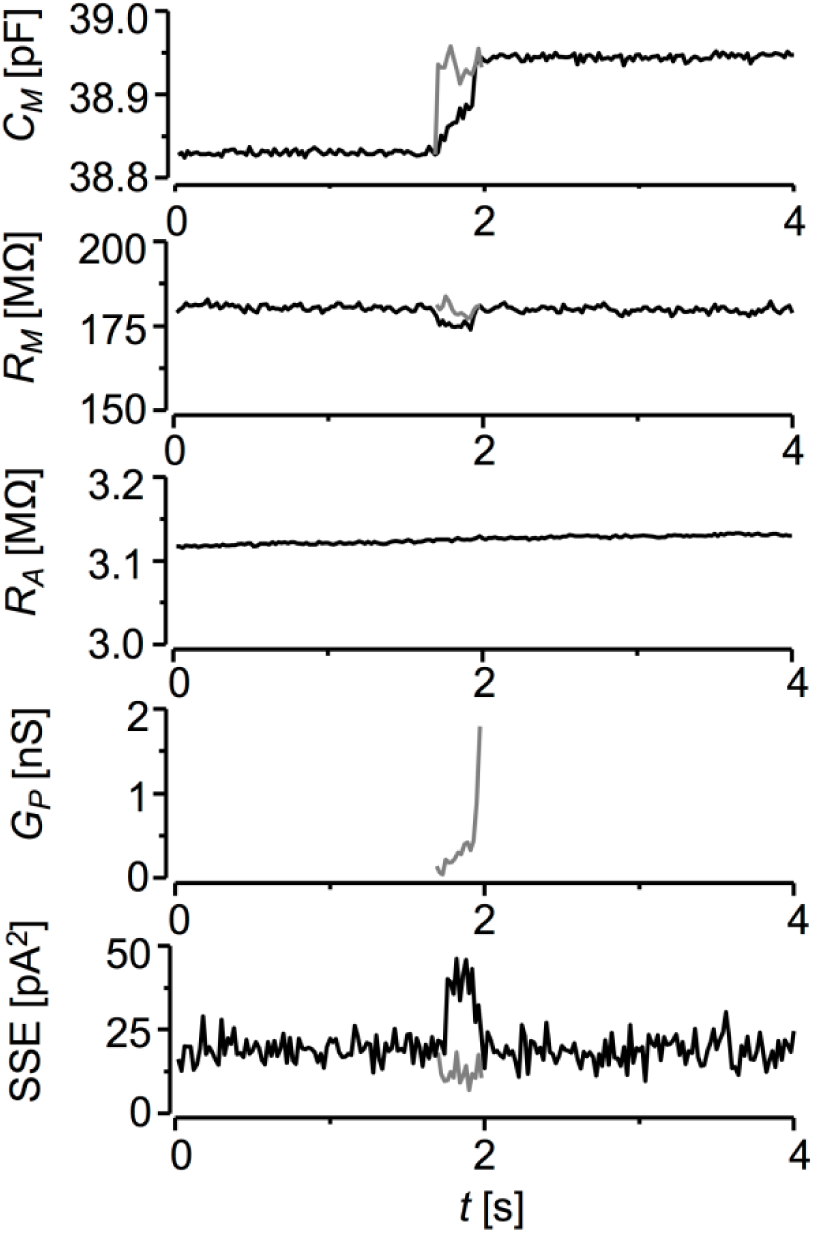
Detection of the fusion pore. A segment of current responses displaying about 100 fF membrane capacitance increase was analysed either by the three-element model (Eq. 1, black traces) or with the five-element model (Eq. 7, grey traces) of the input impedance. Isolated ventricular myocyte of neonatal rat heart, holding potential of 0 mV, square-wave ±40 mV, 1.25 ms period, bandwidth 50 Hz.

### Analysis of seal resistance variation

There is a well-known but also well-hidden problem of the stability of seal resistance that is common to all patch-clamp measurements, since a stable and well isolating seal is needed to confine the circuit current to the cell membrane (Fig 1A). The seal resistance is difficult to measure [18] but its quality can be assessed indirectly [15,19]. If cell membrane resistance reaches the gigaohm range, a small variation of *R*_*Seal*_ may cause the recorded current to be at variance with the membrane current. Thus, changes in *R*_*Seal*_ may affect estimates of current parameters in impedance measurements and generate artefactual events in current records. We tested the effect of *R*_*Seal*_ by simulation of the whole circuit (Fig 1A) using Eq. 5. As evidenced in Fig 8A (left panel), reduction of *R*_*Seal*_ below 10 GΩ strongly affected estimates of *C*_*M*_ and *R*_*M*_ but had negligible effect on *R*_*A*_ estimates. Importantly, however, it had no effect at all on the charge under the capacitive membrane current response *Q*_*C*_, calculated as a simple integral of the current response above the steady current level according to Eq. 8:

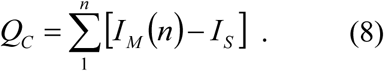

**Fig 8.**
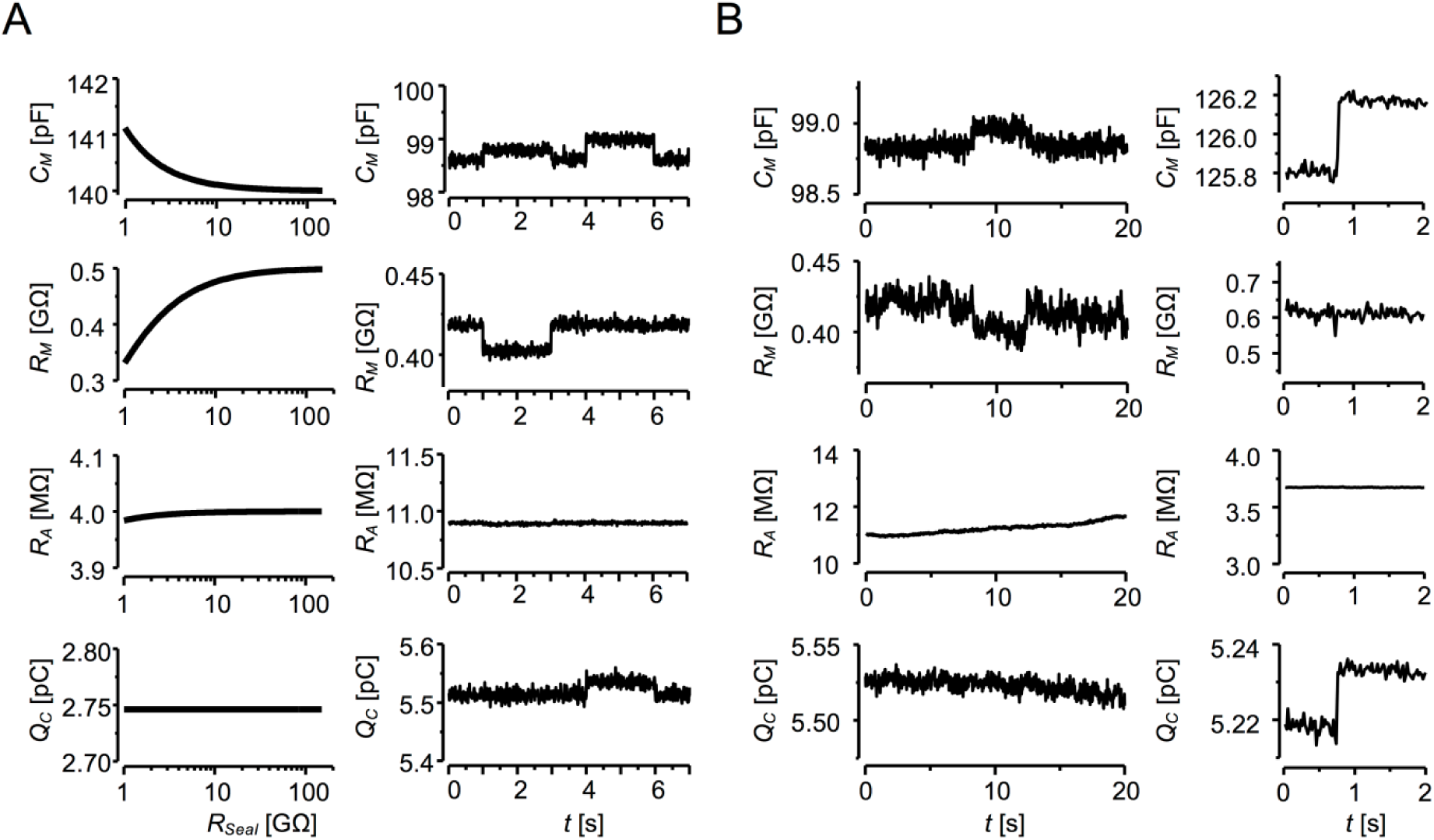
Effect of the seal resistance on circuit parameters. **(A)** Simulated data. *Left panel*: Dependence of estimates of cell impedance parameters on *R*_*Seal*_ value (estimated by Eqs. 1 and 8 from simulated current responses of the equivalent circuit according Eq. 5, Fig 1A, *C*_*M*_ = 140 fF, *R*_*M*_ = 0.5 GΩ, *R*_*A*_ = 4 MΩ). *Right panel*: Simulated experiment with two events. The first event contained a step decrease of *R*_*Seal*_ from 100 to 10 GΩ while the second event contained a step change of *C*_*M*_ from 98.7 fF to 99.0 fF. Note that in the first event an anti-correlated step change emerged in *C*_*M*_ and *R*_*M*_ but not in *R*_*A*_ or *Q*_*C*_ traces, while in the second event a correlated change was present in the *C*_*M*_ and *Q*_*C*_ but not in *R*_*M*_ and *R*_*A*_ traces. The same set of circuit parameters as in the left panel, square-wave ±30 mV, 20 ms period, analysed by the deconvolution procedure. **(B)** Records of impedance parameters of ventricular myocytes obtained by the deconvolution procedure. *Left panel*: A segment of a record containing an anti-correlated step change in *C*_*M*_ and *R*_*M*_ but not in *R*_*A*_ or *Q*_*C*_ traces (square-wave ±30 mV, 20 ms period), indicating artefactual change in membrane capacitance arising from the change of *R*_*Seal*_. *Right panel*: A segment of a record containing a well-resolved step *C*_*M*_ event without correlated changes in either *R*_*M*_ or *R*_*A*_ but mirrored in *Q*_*C*_ (square-wave ±20 mV, 10 ms period), indicating a true change of membrane capacitance.

The illustrative experiment on a cardiomyocyte (Fig 8B) shows an upsurge of *C*_*M*_ that correlated with the drop in *R*_*M*_ but not in *R*_*A*_ and *Q*_*C*_ records. Since the deconvolution-based procedure is virtually free of the cross-talk error, the observed combination of events indicates that this change did not happen at the cell membrane and should be considered as an artefact due to variation of *R*_*Seal*_. Indeed, in the case of true *C*_*M*_ change, as shown in Fig 8B, the large event in the *C*_*M*_ record was not correlated with *R*_*M*_ or *R*_*A*_ but was reflected in *Q*_*C*_.

The charge *Q*_*C*_ was unaffected by *R*_*Seal*_ as expected, because only *I*_*M*_ but not *V*_*M*_ is affected by *R*_*Seal*_. This trait is a precious signature of an unstable seal. Independence of *Q*_*C*_ on step change of *R*_*Seal*_ was confirmed in a simulated experiment (Fig 8A, right panel). A step change of *R*_*Seal*_ caused artefactual increase of *C*_*M*_, decrease of *R*_*M*_, and no change of *R*_*A*_, in their respective traces. For comparison, a true increase of *C*_*M*_ caused no change of *R*_*M*_ and *R*_*A*_ but caused a correlated increase in *Q*_*C*_.

## Discussion

The deconvolution procedure presented and tested in this work would be helpful in studies relying on membrane current recordings obtained under whole-cell voltage-clamp conditions, where an antialiasing filter between the output of the amplifier and the input of the digitizer distorts current records and limits the useful bandwidth of recording. This holds especially for such experiments when step voltage pulses are applied to the cell and the membrane current changes are very fast, as is the case of membrane impedance measurements.

The deconvolution method substantially improved performance of the high-resolution membrane capacitance measurement by suppression of the crosstalk error and by increase of absolute accuracy. Other important benefits come from the simplification and standardization of the analysis procedure. As a result, the deconvolution method reports membrane capacitance changes that can be interpreted with confidence and even semi-automatically. The whole procedure, as implemented in the MAT-MECAS software (http://mat-mecas.sourceforge.net), works with cells of diverse size and membrane capacitance and with common patch-clamp setups in a wide range of experimental conditions and designs.

The method deals well with three critical issues common to all patch-clamp based approaches; namely, the parasitic capacitance, the seal resistance, and the access resistance. These should be fairly stable for reliable performance of standard high-resolution methods. The parasitic capacitance of the recording microelectrode does not create problems for the deconvolution method, if stabilized by a hydrophobic resin applied near the tip of the patch pipette. If stabilized but not electronically compensated at the amplifier, or if overcompensated, *C*_*P*_ might introduce a small error in the estimate of *R*_*A*_. However, due to eliminated crosstalk errors, changes of *C*_*P*_ below 1 pF will not have visible influence on *C*_*M*_ records. The issue of the seal resistance can be overcome by thorough formation of the gigaseal contact [18,19]. *R*_*Seal*_ values above 10 GΩ are safe for artefact-free recordings in most situations. If *R*_*Seal*_ loses its stability and drops to the low-GΩ range, it can be recognized as an anti-correlated change in *C*_*M*_ and *R*_*M*_ traces and no change in *R*_*A*_ and *Q*_*C*_ traces.

Although the deconvolution method is insensitive to variation of *R*_*A*_, the *R*_*A*_ value determines the time constant of recording, and so the optimal rate of stimulation, which should be considered for the best *C*_*M*_ resolution [8]. Faster stimulation rates provide higher resolution [13]; however, a short square-wave period provides less data points for fitting, increases the contribution of noise, and decreases accuracy. Therefore, the decision regarding optimal patch pipette resistance for a given cell size is a trade-off between resolution and accuracy.

Variable membrane conductance does not introduce cross-correlations but increases the noise of *C*_*M*_ records. Therefore, measures that minimize membrane conductance changes, such as proper setting of the holding potential and the stimulus amplitude, or the use of specific channel blockers, should be considered under specific experimental conditions.

A critical issue for current reconstruction by deconvolution is the electrical noise picked up by the recording system. Therefore, proper grounding and shielding of the electrophysiological setup is important. Every peak in the frequency characteristics of a system would be augmented upon reconstruction and would cause oscillations in the time domain.

In this study, we presented results obtained for 10-kHz low-pass filtering and 100-kHz sampling rate. The question was, whether an increased bandwidth would further improve the accuracy and resolution of the measurements. To this end we tested the FFT deconvolution procedure with the low-pass filter set to 100 kHz and the digitizer set to 250 kHz. Test measurements on hardware cells did not confirm the expectations, because the noise in current records was substantially increased. Moreover, the much larger volume of data increased the demands on computational time and resources. On the contrary, the deconvolution approach worked very well with 5, 2, and 1 kHz filtering combined with 50 or 10 kHz digitization, respectively (not shown).

The deconvolution procedure increased the bandwidth of the recorded current response; however, the essence of its success was in correct restoration of the time course of current responses. Thus in *C*_*M*_ recording it brought compliance of recorded currents with theoretical models. In the case of mono-exponential current decay, approximation of the reconstructed current response by the theoretical equation became correct and robust for a wide span of current parameters. The minimal time constant that returned reliable results was 20 μs, corresponding to a cell of 2 pF capacitance (8 μm in diameter) measured with 10 MΩ access resistance.

## Acknowledgments

The authors are grateful to J. Púčik for advice on signal processing and analysis, R. Janíček for testing of the method, A. Zahradníková for consultations and comments on manuscript, and G. Gajdošíková for isolation of cardiac myocytes. The work was supported by APVV-0721-10, APVV-15-0302, and VEGA 2/0147/14.

## Authors’ contribution

MH worked on the method development, designed the deconvolution procedure, performed experiments, analysed the data, performed simulations, wrote software MAT-MECAS and participated in writing the paper. IZ designed and supervised the project, wrote the paper, worked on the method development and designed experiments and the software.

